# Non-canonical cell death by reassortant reovirus

**DOI:** 10.1101/2020.05.20.107706

**Authors:** Roxana M. Rodríguez Stewart, Vishnu Raghuram, Jameson T.L. Berry, Bernardo A. Mainou

## Abstract

Triple-negative breast cancer (TNBC) constitutes 12% of all breast cancer and is associated with worse prognosis compared to other subtypes of breast cancer. Current therapies are limited to cytotoxic chemotherapy, radiation, and surgery, leaving a need for targeted therapeutics to improve outcomes for TNBC patients. Mammalian orthoreovirus (reovirus) is a nonenveloped, segmented, dsRNA virus in the *Reoviridae* family. Reovirus preferentially kills transformed cells and is in clinical trials to assess its efficacy against several types of cancer. We previously engineered a reassortant reovirus, r2Reovirus, that infects TNBC cells more efficiently and induces cell death with faster kinetics than parental reoviruses. In this study, we sought to understand the mechanisms by which r2Reovirus induces cell death in TNBC cells. We show that r2Reovirus infection of TNBC cells of a mesenchymal-stem like (MSL) lineage downregulates the MAPK/ERK pathway and induces non-conventional cell death that is caspase dependent, but caspase 3-independent. Infection of different MSL lineage TNBC cells with r2Reovirus results in caspase 3-dependent cell death. We map the enhanced oncolytic properties of r2Reovirus in TNBC to epistatic interactions between the Type 3 Dearing M2 gene segment and Type 1 Lang genes. These findings suggest that the genetic composition of the host cell impacts the mechanism of reovirus-induced cell death in TNBC. Together, our data show that understanding host and virus determinants of cell death can identify novel properties and interactions between host and viral gene products that can be exploited for the development of improved viral oncolytics.

**Importance:** Triple negative breast cancer (TNBC) is unresponsive to hormone therapies, leaving patients afflicted with this disease with limited treatment options. We previously engineered an oncolytic reovirus (r2Reovirus) with enhanced infective and cytotoxic properties in TNBC cells. However, how r2Reovirus promotes TNBC cell death is not known. In this study, we show that reassortant r2Reovirus can promote non-conventional caspase-dependent but caspase 3-independent cell death and that the mechanism of cell death depends on the genetic composition of the host cell. We also map the enhanced oncolytic properties of r2Reovirus in TNBC to interactions between a Type 3 M2 gene segment and Type 1 genes. Our data show that understanding the interplay between the host cell environment and the genetic composition of oncolytic viruses is crucial for the development of efficacious viral oncolytics.

## Introduction

Breast cancer is the leading cause of cancer in women and second leading cause of death by cancer in women in the United States (https://seer.cancer.gov/). Triple-negative breast cancer (TNBC) constitutes 10-15% of breast cancer diagnoses, has a higher rate of relapse, and lower survival after metastasis than other types of breast cancer (1). TNBC is characterized by its lack of expression of estrogen receptor (ER), progesterone receptor (PR), and human epidermal growth factor receptor 2 (HER-2). These characteristics render TNBC cells unresponsive to hormone therapies that have been efficacious in treating other types of breast cancer (2, 3).

Mammalian orthoreovirus (reovirus) is a segmented double-stranded RNA (dsRNA) virus in the *Reoviridae* family (4). Reovirus has three large (L1, L2, L3), three medium (M1, M2, M3), and four small (S1, S2, S3, S4) gene segments that encode 8 structural and 3 non-structural proteins (5, 6). There are three reovirus serotypes (types 1, 2, and 3) determined by the recognition of the S1-enconded s1 attachment protein by neutralizing antibodies (4, 7). In humans, reovirus infection usually occurs during childhood, though infection is generally asymptomatic (4, 8-10). Additionally, reovirus preferentially replicates and kills tumor cells (11-14). Because of these features, a lab adapted type 3 reovirus is currently in Phase I-III clinical trials to test its efficacy against a variety of cancers (https://clinicaltrials.gov). However, little is known about the biology of reovirus infection in TNBC.

TNBC cells are categorized into subtypes based on their genetic composition (1, 15). Cells in the mesenchymal stem-like (MSL) subtype, including MDA-MB-231 and MDA-MB-436 cells, are characterized by enriched expression of genes involved in motility, cellular differentiation, and growth factor pathways (15-23). The K-Ras G13D and B-Raf G464V B-Raf mutations found in MDA-MB-231 cells result in an upregulated Ras pathway (24, 25). Constitutively active Ras mutations have been identified in many human tumors and signaling through Ras increases tumor cell proliferation and survival in some cancers (26-29). B-Raf regulates the Raf–mitogen-activated protein kinase (MAPK)/extracellular signal-regulated kinase (ERK) pathway by phosphorylation of MEK 1/2, which activates the kinase (30). MAPK/ERK signaling promotes cancer cell proliferation, survival, and metastasis (31). Small-molecule inhibitors that target various steps of the MAPK/ERK pathway are currently in clinical trials to test their efficacy against several cancers (32).

Activated Ras signaling regulates various aspects of reovirus biology, including virus uncoating, infectivity, replication, and release from infected cells (13, 14, 33-40). However, reovirus can also infect and kill cancer cells independent of Ras activation (41-44). In some cells, reovirus downregulates Ras signaling during infection, inducing programmed cell death (45). Reovirus can induce cell death by apoptosis, necroptosis, cell cycle arrest or autophagy (36, 46-60). Reovirus can trigger apoptosis through recognition of viral nucleic acid by cellular pattern recognition receptors and subsequent activation of caspase 8, Bid cleavage, and disruption of the mitochondrial membrane. This results in cytochrome c release, caspase 9 activation, and activation of executioner caspases 3 and 7 (4, 46-49, 52, 54, 55, 61-68). Reovirus can also induce caspase-independent cell death through induction of RIPK3 and MLKL-dependent necroptosis (50, 51, 57). The mode of cell death induced by reovirus appears to be largely dependent on the host cell.

We previously engineered an oncolytic reovirus with enhanced infective and cytotoxic properties in TNBC (r2Reovirus) (69). Oncolytic r2Reovirus is a reassortant virus with 9 gene segments from serotype 1 Lang (T1L) reovirus and a serotype 3 Dearing (T3D) M2 gene segment, as well as several synonymous and non-synonymous point mutations. Strain-specific differences in infectivity, replication, and induction of cell death indicate a vital role of specific viral factors in defining the host cell response and outcome of infection (70-72) (46, 55, 63, 64, 73). It is not known how r2Reovirus promotes TNBC cell death or the contribution of specific viral factors to the enhanced oncolytic properties of the virus.

In this study, we sought to better understand how reovirus induces programmed cell death in a subtype of TNBC and the viral factors associated with this phenotype. We show that reassortant r2Reovirus can promote TNBC cell death by inhibiting MAPK/ERK signaling and inducing a non-conventional cell death that is caspase dependent, but caspase 3-independent conditional on the genetic composition of the host cell. These data suggest that the genetic composition of the host cell can greatly impact the type of cell death induced by reovirus. We also show that the enhanced oncolytic properties of r2Reovirus in TNBC likely map to the presence of a T3D M2 gene segment in the context of an otherwise T1L virus. Together, our data show that an improved understanding of host cell and virus interactions can identify biological properties and interactions between viral gene products to better understand how viruses promote cell death and exploited for the development of improved viral oncolytics.

## Results

### r2Reovirus impairs MAPK/ERK signaling

We previously generated a reassortant reovirus with enhanced infective and cytotoxic properties in TNBC cells (69). The mechanisms through which this virus promotes TNBC cell death is not known. It is also largely unclear how reoviruses promote TNBC cell death. The MDA-MB-231 TNBC cell line has an upregulated Ras pathway from mutations in Ras (G13D) and B-Raf (G464V) (24). To determine the effect of parental reoviruses (T1L and T3D) and r2Reovirus on MAPK/ERK, MDA-MB-231 cells were infected with mock, T1L, T3D, or r2Reovirus at an MOI of 500 PFU/cell or treated with 10 μM MEK1/2 inhibitor U0126. Whole cell lysates were collected at 0, 1, and 2 days post-infection (dpi) and probed for phosphorylated and total MEK1/2 and ERK1/2 by immunoblot (Fig. 1A). Infection with T1L and T3D did not affect the levels of phosphorylated MEK1/2 or ERK1/2 when compared to uninfected cells (mock) at the times tested (Fig. 1B). In cells infected with r2Reovirus, levels of phospho- and total MEK1/2 and total ERK1/2 were slightly lower than mock and levels of phospho-ERK1/2 were significantly lower at 2 dpi than mock. These data suggest that infection with r2Reovirus, but not T1L nor T3D, results in downregulation of MAPK/ERK pathway in these cells.

To determine the effect of inhibiting MAPK/ERK signaling on reovirus-infected MDA-MB-231 cell viability, cells were treated with increasing concentrations of U0126 for 1 h, adsorbed with mock, T1L, T3D, or r2Reovirus at an MOI of 100 PFU/cell or 50 μM etoposide as a positive control, and cell viability was assessed over 6 days (Fig. 2A). Similar to that observed previously (69), r2Reovirus impaired cell viability with faster kinetics than T1L and T3D did not impact cell viability. Treatment of cells with U0126 alone resulted in a dose-dependent cytostatic effect on cell viability, with cell viability leveling at 2 dpi when treated with 10 μM U0126 (red line). This was expected as MAPK/ERK signaling is necessary for cell proliferation in these cells (74-80). Treatment with 0.1 μM U0126 had no significant effect on cell viability in the presence or absence of reovirus. Infection of cells with T3D in addition to U0126 did not significantly impact U0126-induced cytotoxicity. Infection with T1L in the presence of U0126 enhanced the cytotoxicity kinetics, with infection in the presence of 5 or 10 μM U0126 having an additive effect on cytotoxicity (Fig. 2B). Similar to that observed with T1L, Infection of cells with r2Reovirus in the presence of U0126 enhanced the kinetics of cytotoxicity, with 10 μM U0126 having a significant combinatorial effect on cytotoxicity induced by the drug and virus alone. These data show that inactivation of MEK-ERK signaling in MDA-MB-231 cells impairs cellular proliferation without having a cytotoxic effect on cells and that in the context of infection with T3D, does not promote viral cell killing. While inactivation of this pathway in the context of infection with T1L or r2Reovirus enhances the cytotoxic effect of the virus, a statistically significant impairment on cell viability was only observed in the presence of r2Reovirus and 10 μM U0126. Together, these data show r2Reovirus downregulates MAPK/ERK signaling and infection with a serotype 1 reovirus in the presence of a MEK inhibitor enhances the kinetics of viral-mediated cytotoxicity in these cells.

**FIG 1.**
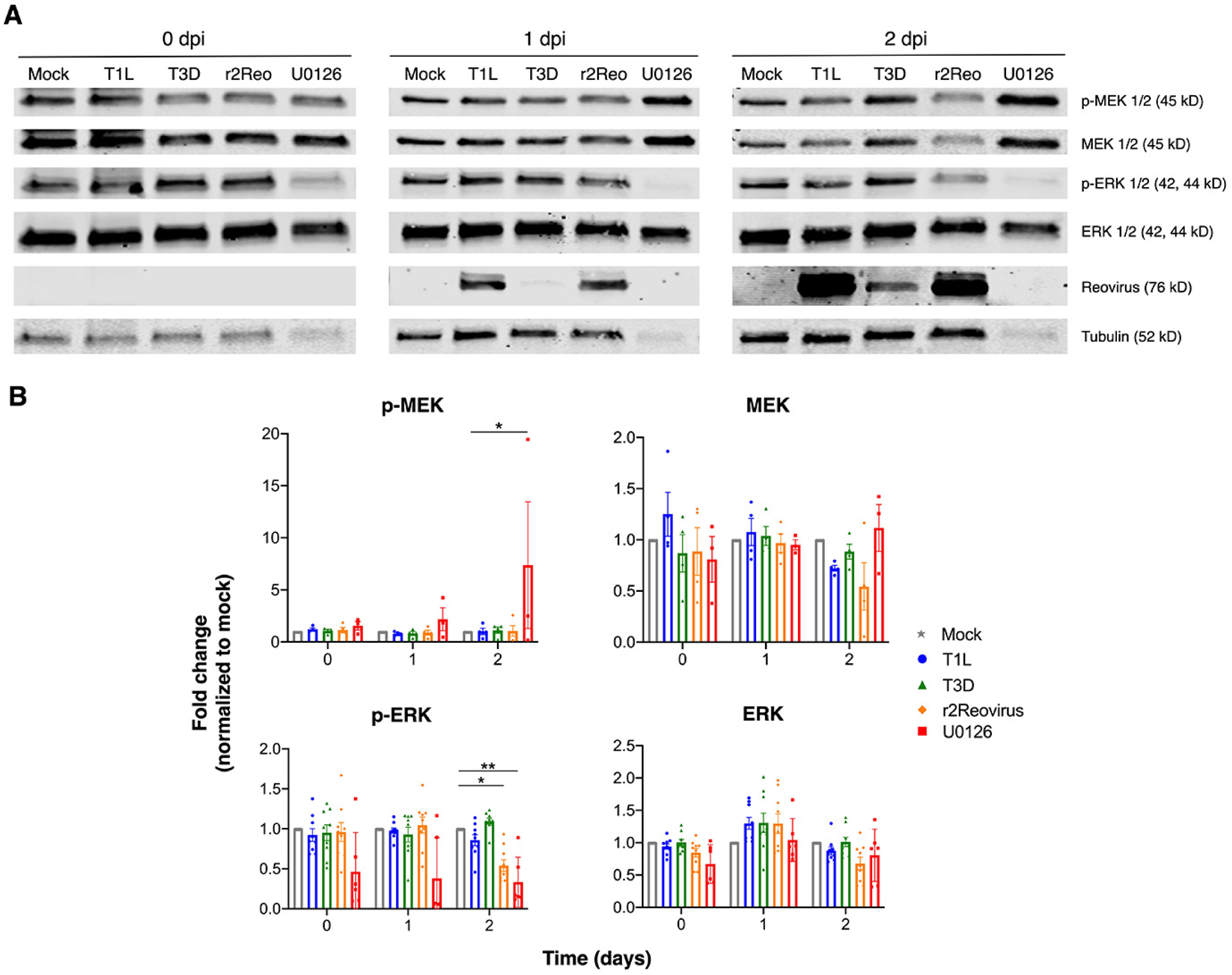
r2Reovirus downregulates MAPK/ERK signaling. MDA-MB-231 cells were adsorbed with mock, T1L, T3D, or r2Reovirus (r2Reo) at an MOI of 500 PFU/cell or treated with 10 μM U0126 for 1 h. A) Whole cell lysates were collected at 0-2 dpi, resolved by SDS-PAGE, and immunoblotted with antibodies specific for phosphorylated and total MEK and ERK, reovirus, and tubulin. Representative data of independent experiments shown. B) Quantitation of band intensity from five independent experiments for phosphorylated MEK (p-MEK) and total MEK and nine independent experiments for phosphorylated ERK (p-ERK) and total ERK and SEM. Data are normalized to mock. *, *P* ≤ 0.002; **, *P* < 0.0001 in comparison to mock at each time point, as determined by two-way ANOVA with Tukey’s multiple-comparison test.

**FIG 2.**
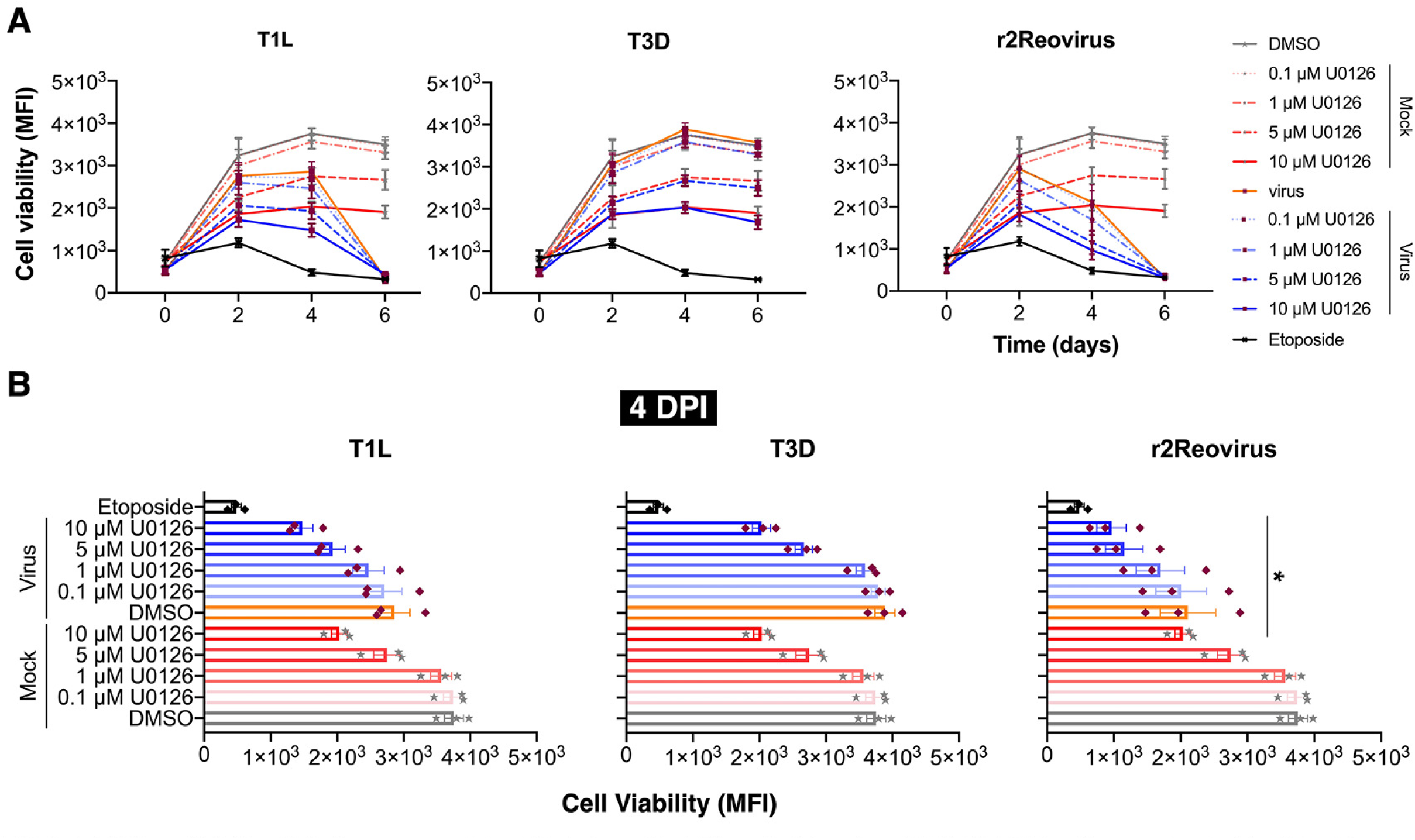
Inhibition of MEK activity increases cytotoxicity induced by T1L and r2Reovirus. MDA-MB-231 cells were treated for 1 h with vehicle (DMSO) or increasing concentrations of U0126 and adsorbed with mock, T1L, T3D or r2Reovirus at an MOI of 100 PFU/ml or treated with 50 μM etoposide for 1 h. A) Cell viability was assessed at times shown. Results are presented as mean fluorescence intensity (MFI) and SEM for three independent experiments. B) Cell viability for data shown in A for day 4 pi. *, *P* ≤ 0.04; **, *P* = 0.001; ***, *P* ≤ 0.005 in comparison to virus alone, as determined by two-way ANOVA with Tukey’s multiple-comparison test.

### Induction of cell death by r2Reovirus is partially dependent on caspases

Inhibition of MAPK/ERK signaling can result in the induction of apoptosis (45, 81-84) and reovirus can induce apoptosis *in vitro* and *in vivo* (4, 46-49, 52, 54, 55, 61-68). To determine if r2Reovirus induces caspase-dependent cell death in TNBC cells, MDA-MB-231 cells were treated with vehicle (DMSO) or 25 μM pan-caspase inhibitor Q-VD-OPH for 1 h, infected with mock or r2Reovirus at an MOI of 500 PFU/cell, and assessed for cell death by annexin V-fluorescein isothiocyanate (FITC)/propidium iodide (PI) staining over 3 days (Fig. 3A). Following r2Reovirus infection, 35.43% of infected cells were annexin V+/PI+ by 2 dpi and 49.24% by 3 dpi. Infection in the presence of Q-VD-OPH decreased the percentage of annexin V+/PI+ cells to 19.09% at 2 dpi and 30.79% by 3 dpi. In etoposide-treated cells, 48.39% of cells were annexin V+/PI+ by day 2 post treatment and 78.12% by day 3 post treatment. Q-VD-OPH treatment decreased the percentage of annexin V+/PI+ to 10% by day 2 post treatment and 20% by day 3 post treatment. Q-VD-OPH also increased the number of annexin V-/PI-cells during infection or etoposide treatment, especially at 2 dpi. Interestingly, we did not observe a significant number of annexin V+/PI-cells under any condition tested. These data show that the pan caspase inhibitor Q-VD-OPH dampens, but does not fully block, r2Reovirus-induced cell death.

**FIG 3.**
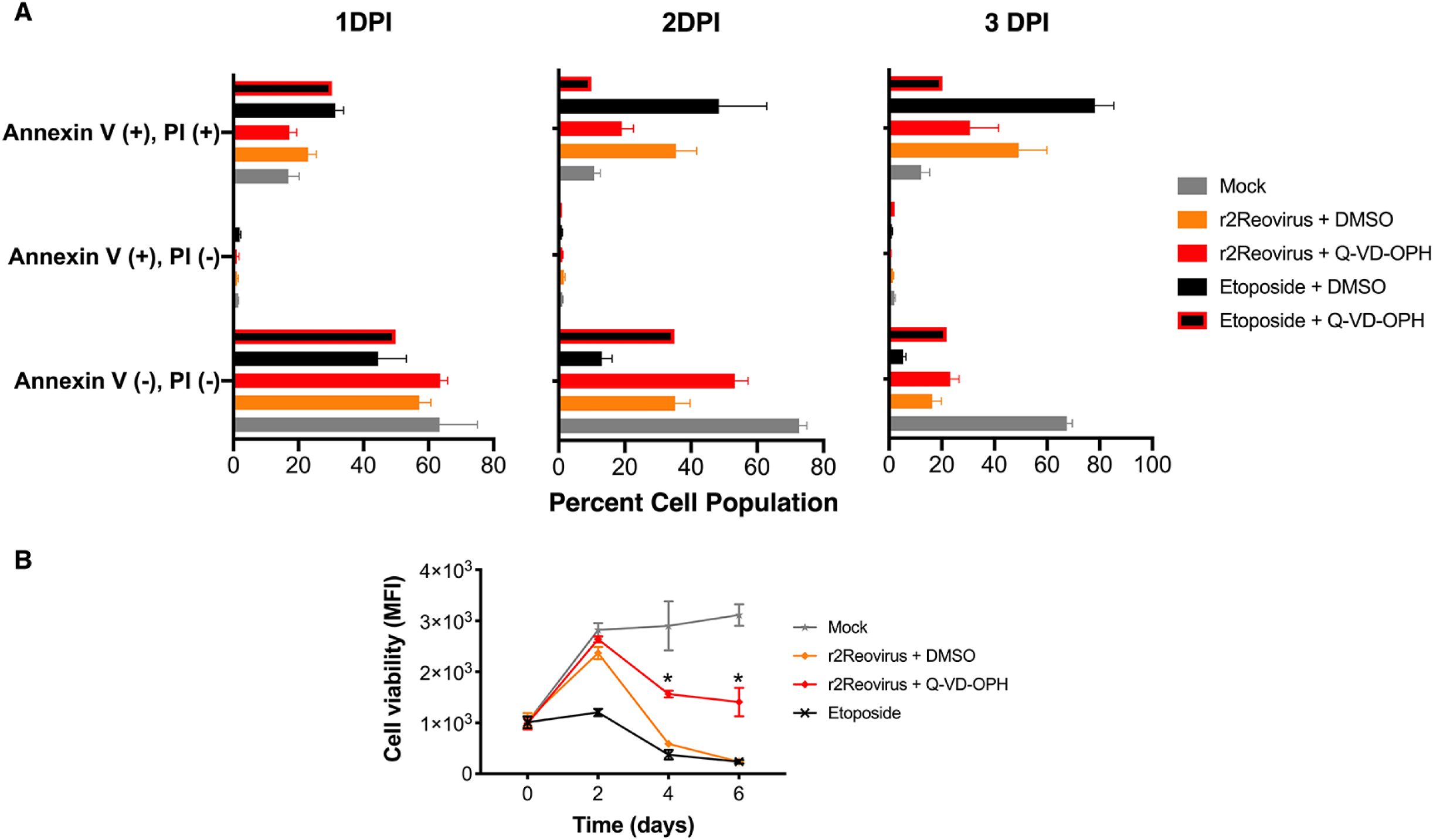
r2Reovirus induced cell death is partially dependent on caspases. MDA-MB-231 cells were treated for 1 h with vehicle (DMSO) or 25 μM caspase inhibitor Q-VD-OPH and adsorbed with mock or r2Reovirus at an MOI of 500 PFU/ml or treated with 50 μM etoposide for 1 h. A) Cells were assessed for annexin V and PI levels by flow cytometry at times shown. Data shown as percentage of cells that are AV-/PI-, AV+/PI- or AV+/PI+. B) Cell viability was assessed at the times shown. Results are shown as mean fluorescence intensity (MFI) and SEM for three independent experiments. *, *P* ≤ 0.0002 in comparison to r2Reovirus, as determined by two-way ANOVA with Tukey’s multiple-comparison test.

To further assess if r2Reovirus is dependent on caspases to promote cell death, MDA-MB-231 cells were treated with DMSO or 25 μM Q-VD-OPH for 1 h, infected with mock or r2Reovirus at an MOI of 500 PFU/cell or treated with 50 μM etoposide, and cell viability was assessed over 6 days (Fig. 3B). Infection of cells with r2Reovirus in the presence of Q-VD-OPH significantly impacted viral-induced cytotoxicity at 4 and 6 dpi, with cell viability over two times greater compared to infection in the absence of the drug. These data show that inhibition of caspase activity results in a significant, but not total, reduction of viral-mediated cell death.

### Reovirus does not affect mitochondrial permeability during infection of MDA-MB-231 cells

Reovirus can induce extrinsic and intrinsic apoptosis (4, 46-49, 52, 54, 55, 61-68). During intrinsic apoptosis, mitochondrial membrane permeabilization leads to loss of mitochondrial transmembrane potential, and release of cytochrome c into the cytoplasm (68, 85-88). To assess if reovirus infection of MDA-MB-231 cells impacts mitochondrial membrane potential, cells were infected with mock, T1L, T3D or r2Reovirus at an MOI of 500 PFU/cell or treated with DMSO or 50 μM etoposide and analyzed by flow cytometry over a 3 day time course of infection using tetramethylrhodamine, ethyl ester (TMRE), a positively-charged dye that accumulates in the mitochondria (Fig. 4A). Infection with T1L or T3D did not significantly affect mitochondrial membrane potential at any of the times tested, while infection with r2Reovirus slightly reduced mitochondrial membrane potential at 3 dpi. Treatment with etoposide significantly reduced mitochondrial membrane permeabilization at all times tested, with stark changes at days 2 and 3 post treatment. These results suggest reovirus induces cell death in MDA-MB-231 cells without major disruption of the mitochondrial membrane.

**FIG 4.**
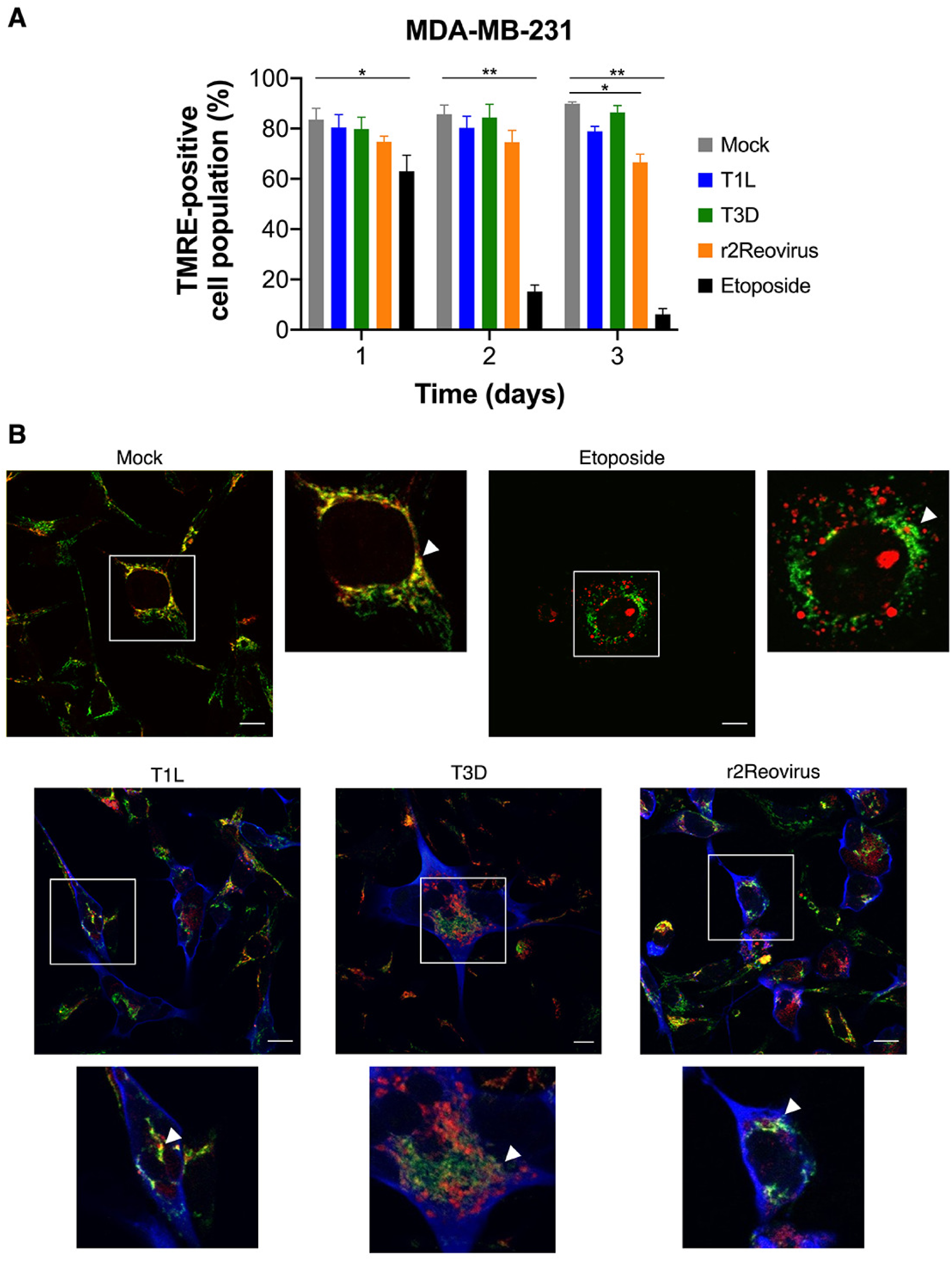
Reovirus does not affect the mitochondria during infection of MDA-MB-231 cells. MDA-MB-231 cells were adsorbed with mock, T1L, T3D, and r2Reovirus for 1 h at an MOI of 500 PFU/cell or treated with 50 μM etoposide. A) Cells were assessed for levels of tetramethylrhodamine, ethyl ester (TMRE) by flow cytometry at times shown. Results are presented as the percentage of TMRE-positive cells and SEM for three independent experiments. *, *P* ≤ 0.008; **, *P* < 0.0001. B) Cells were fixed at 3 dpi and stained with antibodies specific for reovirus (blue) or cytochrome c (green) or with MitoTracker to visualize the mitochondria (red). Images are representative of two independent experiments. Scale bar, 10 μm.

Cytochrome c release from the mitochondria is a key event that can lead to apoptosome formation and subsequent caspase 3 activation. It is possible for cytochrome c to be released from the mitochondria without impacting mitochondrial membrane integrity (89-91). To determine if cytochrome c is released following reovirus infection of MDA-MB-231 cells, cells were infected with mock, T1L, T3D, or r2Reovirus at an MOI of 500 PFU/ml or treated with 50 μM etoposide and assessed for intracellular localization of cytochrome c (green), mitochondria (red), or reovirus antigen (blue) by confocal microscopy at 3 dpi (Fig. 4B). In uninfected cells, cytochrome c largely co-localized with mitochondria. In etoposide-treated cells, cytochrome c localized to areas surrounding swollen mitochondria. In cells infected with T1L, T3D, and r2Reovirus, cytochrome c largely co-localized with mitochondria with no observable swollen mitochondria. These data indicate that during reovirus infection of MDA-MB-231 cells, cytochrome c remains largely associated with mitochondria. Together with TMRE data, these results show that reovirus induces MDA-MB-231 cell death independent of disruption of mitochondrial membrane potential and cytochrome c release.

### Infection with serotype 1 reoviruses increases caspase 9 activity

We next assessed the activation status of caspase 9, a component of the apoptosome that can be activated independent of cytochrome c release in a caspase 8-dependent manner (62, 92). MDA-MB-231 cells were adsorbed with mock, T1L, T3D or r2Reovirus at an MOI of 500 PFU/cell or treated with DMSO or 50 μM etoposide for 1 h, and assessed for caspase 9 activity over a 3 day time course of infection (Fig. 5A). Infection with T1L and r2Reovirus significantly induced caspase 9 activation, with caspase 9 activity levels increasing by 2 dpi and reaching up to 2-fold over mock by 3 dpi. Infection with T3D and treatment with etoposide did not impact caspase 9 activation at the times tested. To measure the requirement of caspase 9 activity in r2Reovirus-mediated cell death of MDA-MB-231 cells, cells were treated with DMSO, 25 μM caspase 9 inhibitor z-LEHD-fmk, or 25 μM Q-VD-OPH for 1 h, infected with mock or r2Reovirus at an MOI of 500 PFU/cell or treated with 10 μM doxorubicin, and assessed for cell viability over 6 days (Fig. 5B). As observed previously, treatment of cells with the pan-caspase inhibitor (Q-VD-OPH) partially blocked reovirus-induced cytotoxicity. Treatment of cells with the caspase 9 inhibitor (z-LEHD-fmk) reduced virus-induced cytotoxicity, albeit not to the same extent as the pan-caspase inhibitor. Together these data show that although infection with T1L and r2Reovirus does not affect mitochondrial membrane potential or promote release of cytochrome c, infection promotes caspase 9 activation. These data also show that although activation of caspase 9 is not solely responsible for viral-mediated cytotoxicity in MDA-MB-231 cells, caspase 9 activation is necessary for the full cytotoxic effects of serotype 1 infection in these cells.

**FIG 5.**
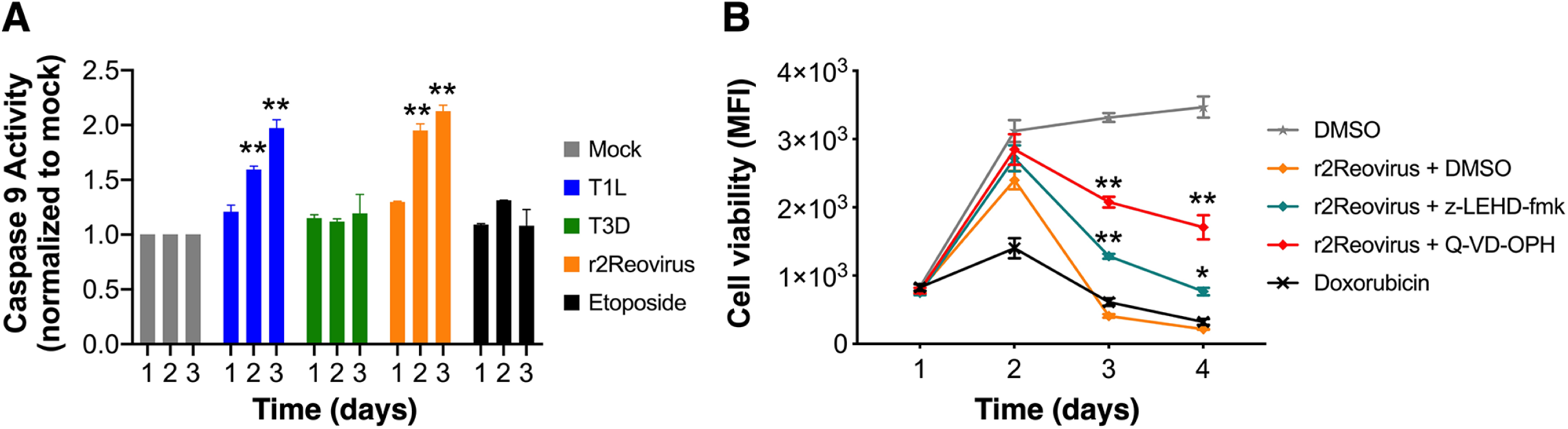
Caspase 9 is activated but not necessary for reovirus-mediated cell death. A) MDA-MB-231 cells were infected with mock, T1L, T3D or r2Reovirus at an MOI of 500 PFU/cell or treated with 50 μM etoposide for 1 h. Cells were assessed for caspase 9 activity at times shown. Results are shown for caspase 9 activity normalized to mock at each time point and SEM for three independent experiments. B) MDA-MB-231 cells were treated with vehicle (DMSO), 25 μM caspase 9 inhibitor z-LEHD-fmk or 25 μM pan-caspase inhibitor Q-VD-OPH and adsorbed with mock or r2Reovirus at an MOI of 500 PFU/cell or treated with 10 μM doxorubicin for 1 h. Cell viability was assessed at times shown. Results are presented as mean fluorescence intensity (MFI) and SEM for three independent experiments. *, *P* = 0.003; **, *P* < 0.0001 in comparison to A) mock and B) r2Reovirus, as determined by two-way ANOVA with Tukey’s multiple-comparison test.

### r2Reovirus blocks caspase 3/7 activity in a replication-dependent manner

Caspase 9 activates caspase 3 and 7 during intrinsic apoptosis (93-95). To determine if caspase 3 is activated during reovirus infection of MDA-MB-231 cells, cells were infected with mock, T1L, T3D or r2Reovirus at an MOI of 500 PFU/cell or treated with 50 μM etoposide, and caspase 3/7 activity was assessed over a 3 day time course of infection (Fig. 6A). Caspase 3/7 activity was not observed in MDA-MB-231 cells infected with mock, T1L, T3D, or r2Reovirus during the times tested. Caspase 3/7 activity was also not observed in cells infected with T1L, T3D, or r2Reovirus at days 4-6 post-infection (data not shown). Treatment of cells with etoposide induced robust caspase 3/7 activity by day 2 post treatment. These data show that etoposide can activate caspase 3 in MDA-MB-231 cells, although in a caspase 9-independent manner. Interestingly, infection with T1L or r2Reovirus does not lead to activation of caspase 3 despite robust caspase 9 activation. These data also show that in MDA-MB-231 cells caspase 3 can be activated.

**FIG 6.**
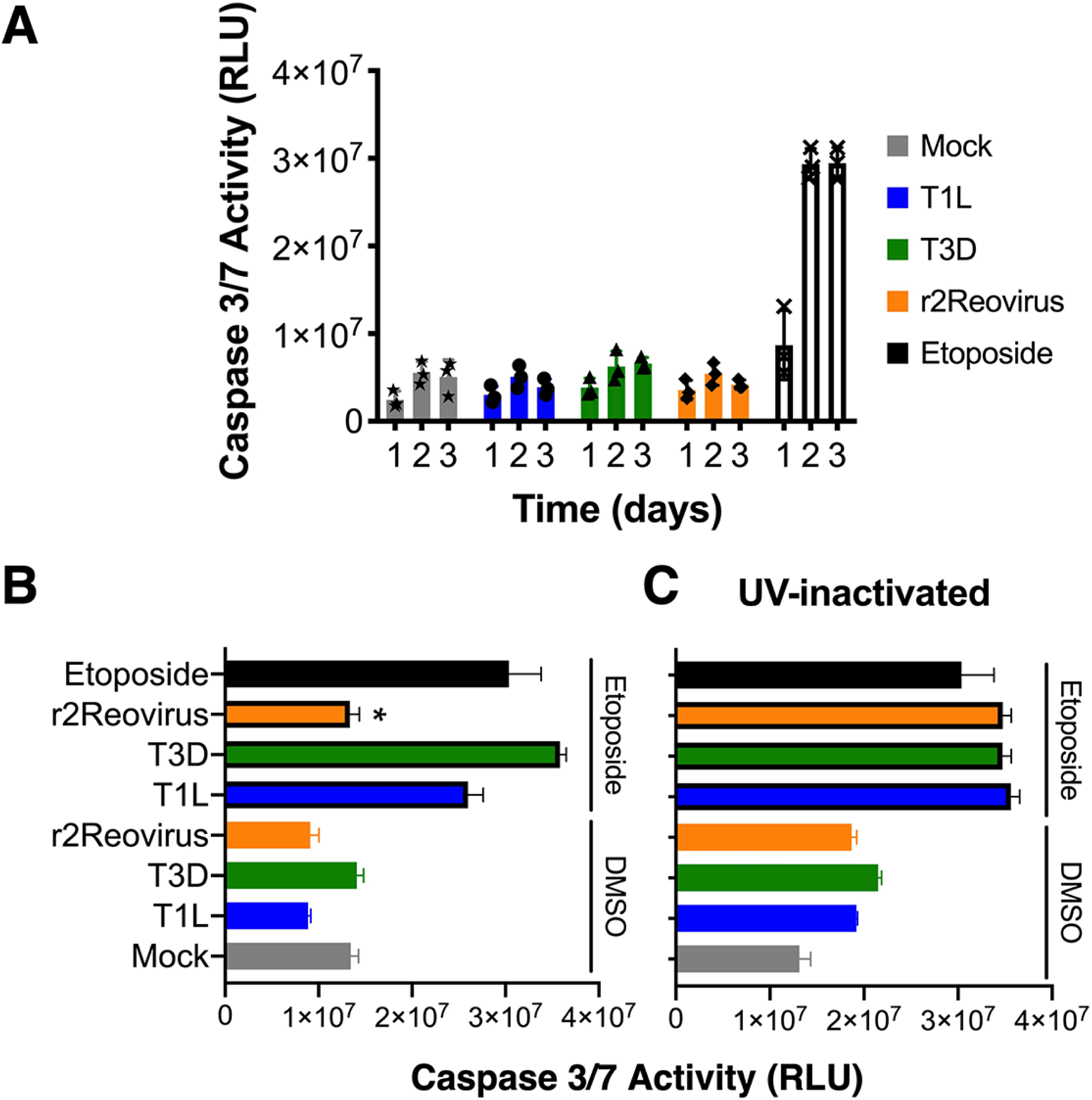
r2Reovirus blocks etoposide-induced caspase 3/7 activity in a replication-dependent manner. A) MDA-MB-231 cells were infected with mock, T1L, T3D or r2Reovirus at an MOI of 500 PFU/cell or treated with 50 μM etoposide for 1 h. Caspase 3/7 activity was measured at times shown in relative luminometer units (RLU) and SEM for three independent experiments. Cells were infected with mock, B) untreated or C) UV-inactivated reovirus at an MOI of 500 PFU/cell, treated with DMSO or 50 μM etoposide 1 dpi, and assessed for caspase 3/7 activity 3 dpi in relative luminometer units (RLU) and SEM for three independent experiments. *, *P* < 0.0001 in comparison to etoposide, as determined by one-way ANOVA with Dunnett’s multiple-comparison test.

To determine if reovirus can impact the activation of caspases 3 and 7, MDA-MB-231 cells were infected with mock, T1L, T3D, or r2Reovirus for 1 day, treated with DMSO or 50 μM etoposide, and assessed for caspase 3/7 activity 2 days post etoposide treatment (Fig. 6B). As previously seen, infection with reovirus did not induce caspase 3/7 activity, whereas etoposide treatment resulted in robust caspase 3/7 activation. Infection of cells with T1L or T3D prior to etoposide treatment did not significantly affect etoposide-induced activation of caspase 3/7. In contrast, infection with r2Reovirus prior to etoposide treatment fully blocked etoposide-induced caspase 3/7 activation. To determine if the ability of r2Reovirus to impair etoposide-induced caspase 3/7 activation is dependent on viral replication, MDA-MB-231 cells were infected with mock or UV-inactivated T1L, T3D, or r2Reovirus one day prior to etoposide treatment, and caspase 3/7 activity was assessed 2 days post etoposide treatment (Fig. 6C). In contrast to that observed with replicating virus, infection of cells with UV-inactivated T1L, T3D, or r2Reovirus did not affect etoposide-induced caspase 3/7 activation. Interestingly, infection with UV-inactivated T1L, T3D, and r2Reovirus resulted in a small, but consistent activation of caspase 3/7 compared to uninfected cells. These results indicate that r2Reovirus blocks etoposide-induced caspase 3/7 activation in MDA-MB-231 cells in a replication-dependent manner.

### PARP-1 cleavage during reovirus infection results in a cleavage fragment that does not correspond to caspase 3 proteolysis

Poly (ADP-ribose) polymerase (PARP-1) is involved in many cellular processes, including DNA repair, genomic stability, and programmed cell death (96-99). During apoptosis, PARP-1 is cleaved into an 89 kDa fragment by caspase 3 (100) (101-103). PARP-1 can also be cleaved by various proteases during non-apoptotic cell death (100). To assess if PARP-1 is cleaved during reovirus-infection of MDA-MB-231 cells, cells were infected with mock, T1L, T3D or r2Reovirus at an MOI of 500 PFU/cell or treated with 50 μM etoposide, and whole cell lysates were collected at 0, 1, and 2 dpi. Lysates were resolved by SDS-PAGE and immunoblotted with antisera specific for PARP-1, reovirus, and tubulin (Fig. 7). Etoposide treatment resulted in an 89 kDa PARP-1 cleavage fragment at day 2 post treatment, consistent with etoposide activation of caspase 3. In contrast, infection with T1L and r2Reovirus resulted in a 70 kDa PARP-1 cleavage fragment while infection with T3D, which does not impair MDA-MB-231 cell viability, did not result in PARP-1 proteolysis. Treatment with caspase 3 inhibitor (Q-VD-OPH) did not reduce PARP-1 cleavage following reovirus infection (data not shown). Calpains, cathepsins, granzyme A and B, and matrix metalloprotease 2 (MMP-2) can proteolytically cleave PARP into fragments of the molecular weight observed during reovirus infection (100). These data suggest that during T1L and r2Reovirus infection of MDA-MB-231 cells PARP-1 is cleaved by a protease other than caspase 3.

**FIG 7.**
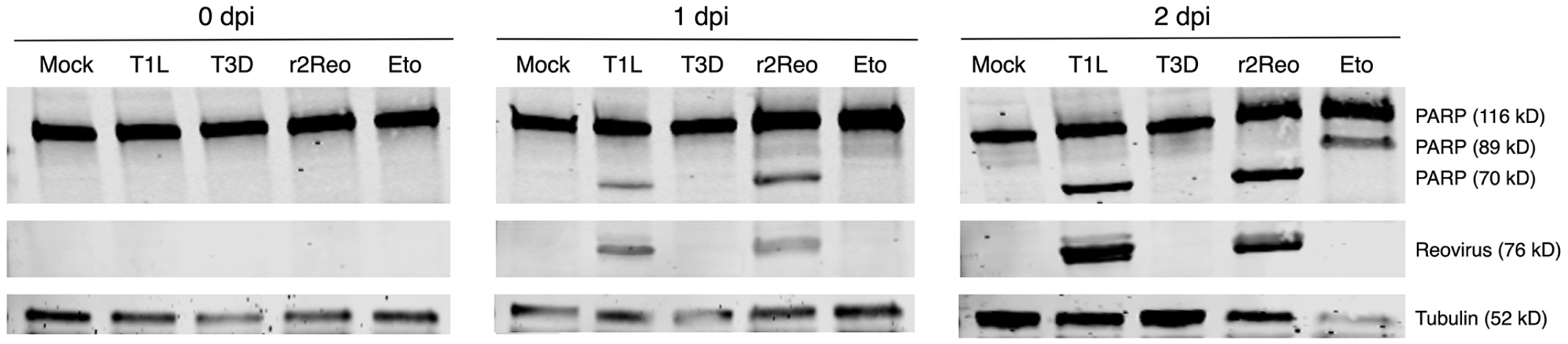
Differential PARP cleavage in reovirus-infected cells. MDA-MB-231 cells were adsorbed with mock, T1L, T3D, or r2Reovirus (r2Reo) at an MOI of 500 PFU/cell or treated with 50 μM etoposide (eto) for 1 h. Whole cell lysates were collected at 0-2 dpi, resolved by SDS-PAGE, and immunoblotted with antibodies specific for PARP, reovirus, and tubulin. Representative experiment of nineteen.

### Reovirus infection of MDA-MB-436 cells promotes caspase 3/7 activation and cell death

r2Reovirus induces cell death with enhanced kinetics in TNBC cells, including MDA-MB-436 cells, a mesenchymal-stem like (MSL) subtype TNBC cell line with different mutations than MDA-MB-231 cells (15, 69). To assess if host heterogeneity affects the mode of cell death induced by reovirus, r2Reovirus oncolysis was tested in MDA-MB-436 cells. To determine if reovirus cell death induction is caspase dependent, MDA-MB-436 cells were treated with DMSO or 25 μM pan-caspase inhibitor Q-VD-OPH for 1 h, infected with mock or r2Reovirus at an MOI of 500 PFU/cell or treated with 50 μM etoposide, and assessed for cell viability over 6 days (Fig. 8A). Infection with r2Reovirus impaired MDA-MB-436 cell viability, with significant cytotoxicity observed by day 4 pi. Treatment of cells with Q-VD-OPH significantly reduced reovirus-mediated cytotoxicity, although cell viability levels were not fully restored to those observed in mock. To determine if r2Reovirus infection of MDA-MB-436 cells impacts mitochondrial membrane potential, cells were infected with mock or r2Reovirus or treated with DMSO or 50 μM etoposide, and assessed for TMRE levels by flow cytometry over 3 days (Fig. 8B). At 2 and 3 dpi, there is a significant decrease in mitochondrial membrane potential in r2Reovirus-infected cells, and to a lesser degree in etoposide-treated cells. To assess if r2Reovirus infection of MDA-MB-436 cells promotes caspase 3/7 activation, cells were infected with mock or r2Reovirus for at an MOI of 500 PFU/cell or treated with DMSO or 50 μM etoposide, and caspase 3/7 activity was assessed over 3 days (Fig. 8C). In contrast to that observed in MDA-MB-231 cells, r2Reovirus infection induced significant caspase 3/7 activation by 1 dpi, with sustained activation over the times tested. Etoposide induced a slight increase in caspase 3/7 activation, but to a lesser degree than reovirus, mirroring that observed by cell viability and TMRE staining. These results show r2Reovirus infection of MDA-MB-436 cells robustly disrupts the mitochondrial membrane and promotes caspase 3/7 activation. These data also show etoposide is not as effective at inducing cell death in MDA-MB-436 cells compared to MDA-MB-231 cells. Together, these data show that the mechanism of cell death induced by r2Reovirus is host cell context-dependent and independent of the ability of the virus to activate caspase 3.

**FIG 8.**
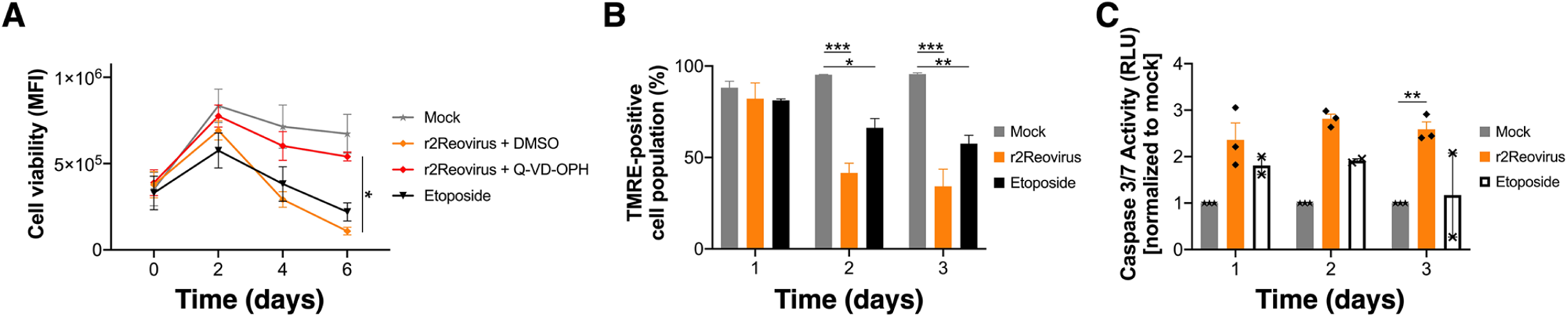
Caspase 3-dependent cell death is observed in MDA-MB-436 cells. A) MDA-MB-436 cells were treated for 1 h with vehicle (DMSO) or 25 μM caspase inhibitor Q-VD-OPH and adsorbed with mock or r2Reovirus at an MOI of 500 PFU/ml for 1 h. Cells were assessed for cell viability over times shown. B, C) MDA-MB-436 cells were infected with mock or r2Reovirus at an MOI of 500 PFU/cell or treated with 50 μM etoposide for 1 h. B) Cells were assessed for TMRE levels by flow cytometry at times shown. Results are presented as the percentage of cells that are TMRE-positive and SEM for three independent experiments. C) Caspase 3/7 activity was measured at times shown. Results are shown as relative luminometer units (RLU) and SEM and normalized to mock for three independent experiments. *, *P* ≤ 0.05; **, *P* ≤ 0.008; ***, *P* < 0.0001 as determined by two-way ANOVA with Tukey’s multiple-comparison test.

### Enhanced r2Reovirus oncolysis maps to the T3D M2 gene segment

The reassortant r2Reovirus is composed of 9 T1L gene segments and an M2 gene segment from T3D in addition to several synonymous and non-synonymous point mutations (69). To determine the contribution of the M2 gene segment to r2Reovirus oncolysis in TNBC cells, reoviruses were engineered by reverse genetics with T1L and T3D M2 gene segment swaps in otherwise isogenic backgrounds (55, 70, 104, 105). To confirm the presence of swapped M2 gene segments, the genetic composition of parental T1L and T3D reoviruses, r2Reovirus, and T1L-T3M2 and T3D-T1M2 was assessed by SDS-gel electrophoresis (Fig. 9A). The electromobility of the reovirus gene segments confirmed that T1L and T3D-T1M2 contain a T1L M2 gene segment, whereas T3D, r2Reovirus, and T1L-T3M2 contain a T3D M2 gene segment (asterisks).

**FIG 9.**
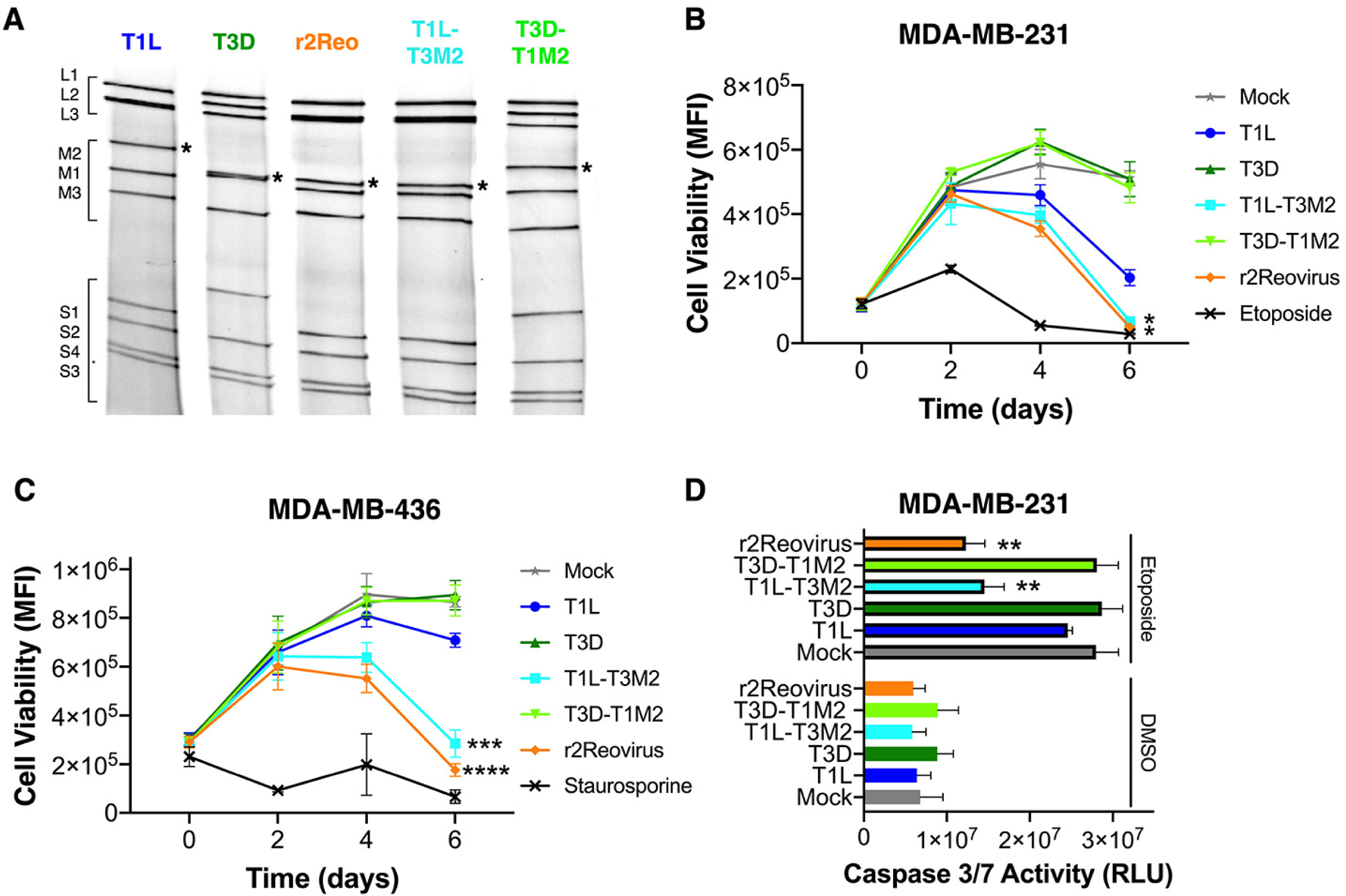
Impact of the M2 gene segment in virus-induced cytotoxicity of TNBC cells. A) SDS-PAGE gel electrophoresis of T1L, T3D, r2Reovirus (r2Reo), and recombinants T1L-T3M2 and T3D-T1M2. Strains are differentiated by migration patterns of three large (L), three medium (M), and four small (S) gene segments. Asterisk denotes M2 gene segment. B) MDA-MB-231 cells were infected with mock, T1L, T3D, T1L-T3M2, T3D-T1M2 or r2Reovirus at an MOI of 500 PFU/cell and treated with vehicle (DMSO) or 50 μM etoposide 1 day post-infection. Caspase 3/7 activity was measured 3 days post-infection. Data are shown as relative luminometer units (RLU) and SEM for three independent experiments. C) MDA-MB-231 and D) MDA-MB-436 cells were infected with mock, T1L, T3D, T1L-T3M2, T3D-T1M2 or r2Reovirus at an MOI of 500 PFU/cell or treated with C) 50 μM etoposide or D) 1 μM staurosporine for 1 h. Cell viability was assessed at times shown. Results are presented as mean fluorescence intensity (MFI) and SEM for three independent experiments. *, *P* ≤ 0.04; **, *P* = 0.009; ***, *P* = 0.0003; ****, P < 0.0001 in comparison to T1L, as determined by two-way ANOVA with Tukey’s multiple-comparison test.

To assess the role of the M2 gene segment in reovirus-induced cytotoxicity of TNBC cells, MDA-MB-231 (Fig. 9B) and MDA-MB-436 cells (Fig. 9C) were infected with mock, T1L, T3D, T1L-T3M2, T3D-T1M2, or r2Reovirus at an MOI of 500 PFU/cell or treated with DMSO, 50 μM etoposide (MDA-MB-231), or 1 μM staurosporine (MDA-MB-436), and assessed for cell viability over 6 days. In both cell lines, T1L-T3M2 impaired cell viability with similar kinetics as r2Reovirus and with faster kinetics than T1L, especially in MDA-MB-436 cells. In contrast, T3D-T1M2 and T3D did not significantly impact cell viability in either MDA-MB-231 or MDA-MB-436 cells. To assess if cytopathic differences observed with the recombinant viruses were a result of differences in infectivity, MDA-MB-231 cells were infected with mock, T1L, T3D, T1L-T3M2 or T3D-T1M2. T1L-T3M2 infected MDA-MB-231 cells at a slightly higher rate than T1L and T3D, while T3D-T1M2 had slightly diminished infectivity when compared to the parental viruses (data not shown). These data indicate that the T3D M2 gene segment is sufficient to enhance the cytotoxic properties of an otherwise T1L virus, without significant impact on infectivity. These data also indicate that the enhanced cytotoxic properties of r2Reovirus map to the T3D M2 gene segment and that mutations found in r2Reovirus likely have little or no effect on the virus’ enhanced oncolysis. Further, the addition of the T1L M2 gene segment to an otherwise T3D virus does not affect the infective or cytopathic properties of the virus. Additionally, even though r2Reovirus induces cell death by different modes in MDA-MB-231 and MDA-MB-436 cells, enhanced cell death induction by the reassortant virus maps to the same viral factor in both cell lines.

To determine if the M2 gene segment impacts the activation of caspase 3/7 following etoposide treatment, MDA-MB-231 cells were infected with mock, T1L, T3D, T1L-T3M2, T3D-T1M2, or r2Reovirus at an MOI of 500 PFU/cell for 1 day, treated with DMSO or 50 μM etoposide, and assessed for caspase 3/7 activity 2 days post etoposide treatment (Fig. 9D). As observed previously, infection with all the reoviruses tested did not induce caspase 3/7 activation and treatment of cells with etoposide resulted in robust activation of caspase 3/7. Infection with T1L-T3M2 or r2Reovirus prior to etoposide treatment robustly impaired caspase 3/7 activation. Conversely, infection with T1L, T3D, or T3D-T1M2 did not impact etoposide-induced caspase 3/7 activation. These results indicate that the ability for r2Reovirus to block etoposide-induced caspase 3/7 activation maps to the T3D M2 gene segment.

## Discussion

Reassortant r2Reovirus infects TNBC cells more efficiently and induces cell death with enhanced kinetics when compared to prototypic strains of reovirus, including the oncolytic reovirus currently in clinical trials. r2Reovirus was obtained from co-infection of MDA-MB-231 cells with T1L, Type 2 Jones (T2J), and T3D followed by serial passaging. r2Reovirus has 9 gene segments from T1L, an M2 gene segment from T3D, and several synonymous and non-synonymous point mutations (69). In this study, we sought to better understand how r2Reovirus, and reoviruses in general, promote TNBC cell death. We focused on MDA-MB-231 cells, a TNBC cell line belonging to the MSL subtype (15). There are limited treatment options against TNBC and the MSL subtype is associated with enrichment of genes involved in cell motility, cellular differentiation, and growth factor signaling pathways (1-3, 15). MDA-MB-231 cells have mutations in *BRAF, CDKN2A, KRAS, NF2, TP53*, and *PDGFRA* genes (15). Mutations in *BRAF* (G464V) and *KRAS* (G13D) result in constitutive activation of MAPK/ERK signaling, promoting survival, proliferation, cell cycle progression, and cell growth (24, 106). Constitutive activation of MAP/ERK is found in several cancers (24, 27-29, 106-109). Constitutive Ras activation can enhance reovirus oncolysis by affecting multiple steps of the viral replication cycle, including enhancing virus uncoating and disassembly, negative regulation of retinoic acid-inducible gene I (RIG-I) signaling and impairing dsRNA-activated protein kinase (PKR) activation, increasing progeny, and enhancing viral spread (35, 40, 110). In some cells, reovirus downregulation of MAP/ERK results in induction of apoptosis (45). In the context of MDA-MB-231 cells, r2Reovirus, but not T1L or T3D, decreased activation of MAPK/ERK signaling. The observed downregulation of MEK 1/2 and ERK 1/2 activation suggests r2Reovirus inhibits this pathway by either directly targeting MEK 1/2 or upstream of MEK 1/2 on Ras and B-Raf, or B-Raf alone. The combination of r2Reovirus and MEK inhibitor U0126 resulted in increased cell death compared to inhibitor or virus alone, highlighting the importance of this pathway in serotype 1 reovirus-mediated cell killing of TNBC cells. The lack of enhancement of cell death by U0126 when combined with T3D suggests that downregulation of MAPK/ERK signaling is not sufficient to promote virus killing.

Downregulation of MAPK/ERK signaling can lead to apoptosis and reovirus can induce programmed cell death by intrinsic and extrinsic apoptosis or necroptosis (36, 45-57). We did not observe an effect on r2Reovirus-induced cell death in the presence of a RIPK3 inhibitor (data not shown), suggesting that reovirus does not promote TNBC cell death by necroptosis. Infection of MDA-MB-231 and MDA-MB-436 cells in the presence of a pan-caspase inhibitor resulted in a significant, but not complete, reduction of virus-mediated cytotoxicity, indicating the need for caspases to promote cell death. During reovirus-induced apoptosis, caspase 8-cleaved Bid translocates to mitochondria, cytochrome c is released following mitochondrial membrane permeabilization, resulting in caspase 9 and caspase 3 activation (4, 46-49, 52, 54, 55, 61-68). In MDA-MB-231 cells, etoposide treatment disrupted mitochondrial membrane potential and promoted cytochrome c release from mitochondria. In reovirus-infected MDA-MB-231 cells, the mitochondrial membrane was largely unaffected and cytochrome c was not released. Despite the lack of disruption of the mitochondrial membrane during reovirus infection, caspase 9 was significantly activated during infection with T1L and r2Reovirus. Caspase 9 can be activated in a cytochrome c-independent manner via caspase 8-activation of caspase 3, which in turn cleaves and activates caspase 9 (62, 92). These results indicate that at least in the context of MDA-MB-231 cells, caspase 9 activation is a property of serotype 1 reoviruses, but not serotype 3 reoviruses. The lack of activation of caspase 3 during reovirus infection suggests a novel mechanism for caspase 9 activation, with the possibility of a viral protein directly activating caspase 9. These findings suggest that in MDA-MB-231 cells, reovirus promotes programmed cell death through a non-canonical pathway.

Activation of caspase 9 can subsequently activate caspase 3 and caspase 7 (93-95). In reovirus-infected MDA-MB-231 cells, caspase 3/7 activity was not observed. Reovirus infection can result in secretion of tumor necrosis factor (TNF)-associated death-inducing ligand (TRAIL), activation of nuclear factor kappa-light-chain-enhancer of activated B cells (NF-κB), and induction of apoptosis (47, 56, 61). There are conflicting data on secretion and sensitivity to TRAIL in MDA-MB-231 cells (111-116). It is possible the lack of effects on the mitochondrial membrane during reovirus infection are linked to MDA-MB-231 cells being insensitive to TRAIL. However, we also observed that r2Reovirus blocks caspase 3/7 activation following etoposide treatment and that this effect is dependent on viral replication.

We show that the ability of r2Reovirus to block caspase 3/7 activation maps to the T3D M2 gene segment. Interestingly, this phenotype is only observed when the T3D M2 gene segment is expressed in the context of T1L, as infection with T3D did not block caspase 3/7 activation. This suggests an epistatic effect of T3D M2 with a T1L-encoded gene. Expression of the T3D-M2 gene segment in the context of an otherwise T1L virus also promoted cell death of MDA-MB-231 and MDA-MB-436 cells with similar kinetics as r2Reovirus and faster kinetics compared to T1L. These data suggest the enhanced cytopathic effects of r2Reovirus in TNBC cells is largely linked to the expression of the T3D M2 gene segment in the context of an otherwise T1L virus, with the point mutations present in r2Reovirus having little or no effect on enhanced oncolysis. T1L-T3M2 reovirus has enhanced attachment and infectivity in L929 and HeLa cells likely due to an interaction between the T3D M2 gene encoded μ1 protein and the T1L attachment protein σ1 (105). In various cells, the S1 and M2 genes are also key factors in reovirus-induced inhibition of cellular DNA synthesis and programmed cell death (46, 63))(49, 64, 117). Though not significant, T1L-T3M2 infected MDA-MB-231 cells at a slightly higher rate than T1L, while T3D-T1M2 showed diminished infectivity when compared to T3D (data not shown). It is possible that in TNBC cells, the interaction between T3D μ1 and T1L *σ*1 promote enhanced oncolysis. These findings further highlight the epistatic effects of reovirus genes in various aspects of reovirus biology.

Several viruses exploit host cell caspases, including caspase 3, to promote viral replication (118-121). Avian reovirus utilizes caspase 3 to proteolytically process viral nonstructural protein μNS, which is involved in the formation of viral factories (122). Mammalian reovirus recruits host proteins to viral factories, including cytoskeletal elements, cellular chaperones, intrinsic immune system proteins, as well as the endoplasmic reticulum (ER) and ER-Golgi intermediate compartment (123-127). It is possible reovirus recruits caspase 3 to viral factories to aid in a step in the replication cycle. Viral protein synthesis and expression of μ1 is required to block necroptosis in L929 cells (57). While we did not observe induction of necroptosis in TNBC cells (data not shown), it is possible that newly synthesized μ1 in conjunction with a T1L gene product blocks caspase 3/7 activation, which results in unconventional cell death in these cells.

During programmed cell death, PARP-1 can be proteolytically cleaved by various proteases (100). As such, PARP-1 cleavage is commonly used as a downstream marker of programmed cell death. Etoposide treatment of MDA-MB-231 cells resulted in an 89 kDa cleaved PARP-1 fragment that is characteristic of caspase 3 cleavage during apoptotic cell death (100, 128) (101-103). Infection with T3D did not result in PARP-1 proteolytic cleavage, which concurs with T3D not promoting cytopathic effects during infection of MDA-MB-231 cells. Infection with T1L or r2Reovirus resulted in a 70 kDa cleaved PARP-1 fragment. Proteases, including calpains, cathepsins, E64-d, Granzyme A and B, and MMP-2 can proteolytically cleave PARP into cleavage fragments of this molecular weight (100). Infection in the presence of a calpain inhibitor blocked PARP-1 cleavage, but the calpain inhibitor also blocked infectivity (data not shown). In addition, treatment with caspase 3 inhibitor did not result in reduced PARP-1 cleavage (data not shown). These results suggest proteolysis of PARP-1 during T1L and r2Reovirus infection is mediated by an enzyme other than caspase 3. These data further suggest that serotype 1 reoviruses promote caspase 3-independent programmed cell death in MDA-MB-231 cells.

The mechanism by which r2Reovirus promotes cell death of another TNBC cell line, MDA-MB-436, was assessed to better understand how host cell heterogeneity impacts virus induced cell death. MDA-MB-436 cells have mutated *BRCA1* and *TP53* genes, and BRAF and KRAS do not have mutations, creating a different cellular environment compared to MDA-MB-231 cells (15). In contrast to that observed in MDA-MB-231 cells, r2Reovirus infection of MDA-MB-436 cells decreases mitochondrial membrane potential and increases caspase 3/7 activity, suggestive of canonical apoptosis. These data suggest that although both TNBC cell lines are more susceptible to r2Reovirus-mediated cell death than parental reoviruses, infection promotes different types of cell death in each cell line. These results suggest that the host cell environment plays a key role in the type of cell death promoted following reovirus infection and that the type of cell death induced by the virus can be independent of the viral genetic composition.

In conclusion, this study identifies a non-conventional virus-induced cell death mechanism in TNBC cells driven by a reassortant oncolytic reovirus. We further map the enhanced cytopathic properties of this reassortant reovirus in TNBC cells to an epistatic effect of the T3D M2 gene with a T1L viral gene product. Better understanding of the interplay between the genetic composition of oncolytic viruses and the host cell environment is crucial for the development of improved reoviruses for oncolytic therapy.

## Materials and Methods

### Cells, viruses, and antibodies

MDA-MB-231 cells (gift from Jennifer Pietenpol, Vanderbilt University) and MDA-MB-436 cells (ATCC^®^ HTB-130™) were grown in Dulbecco’s Modified Eagle’s Medium (DMEM) supplemented with 10% fetal bovine serum (FBS) (Life Technologies), 100 U per ml penicillin and streptomycin (Life Technologies). Spinner-adapted L929 cells (gift from Terry Dermody, University of Pittsburgh) were grown in Joklik’s modified minimal essential medium (MEM) with 5% FBS, 2 mM L-glutamine (Life Technologies), penicillin and streptomycin, and 0.25 mg per ml amphotericin B (Life Technologies).

Reovirus strains Type 1 Lang (T1L) and Type 3 Dearing (T3D) working stocks were prepared following rescue with reovirus cDNAs in BHK-T7 cells (gift from Terry Dermody, University of Pittsburgh), followed by plaque purification, and passage in L929 cells (129). r2Reovirus is a reassortant strain obtained from co-infection of MDA-MB-231 cells with T1L, T2J, and T3D reovirus strains followed by serial passage in these cells. (69). T1L-T3M2 and T3D-T1M2 (gift from Pranav Danthi, Indiana University (130)) were obtained through reovirus reverse genetics (129). Purified virions were prepared using second-passage L929 cell lysate stocks. Virus was purified from infected cell lysates by Vertrel XF (TMC Industries Inc.) extraction and CsCl gradient centrifugation as described (131). The band corresponding to the density of reovirus particles (1.36 g/cm^3^) was collected and dialyzed exhaustively against virion storage buffer (150 mM NaCl, 15 mM MgCl2, 10 mM Tris-HCl [pH 7.4]). Reovirus particle concentration was determined from the equivalence of 1 unit of optical density at 260 nm to 2.1×10^12^ particles (132). Viral titers were determined by plaque assay using L929 cells (133, 134).

Reovirus polyclonal rabbit antiserum raised against reovirus strains T1L and T3D was purified as described (135) and cross-adsorbed for MDA-MB-231 cells. Secondary IRDye 680 and 800 antibodies (LI-COR Biosciences) and goat anti-rabbit Alexa Fluor 405 (A405) (Life Technologies).

### Immunoblotting to assess activation of MAPK/ERK signaling and proteins involved in apoptosis pathway

MDA-MB-231 cells and MDA-MB-436 cells were adsorbed with T1L, T3D, and r2Reovirus at an MOI of 500 PFU/cell for 1 h at room temperature or treated with DMSO or 10 μM U0126, washed with PBS, and incubated for 0-2 days at 37°C. Whole cell lysates were prepared using RIPA buffer (20 mM Tris-HCl [pH 7.5], 150 mM NaCl, 1 mM EDTA, 1% NP-40, 0.1% sodium dodecyl sulfate, 0.1% sodium deoxycholate) and fresh Protease Inhibitor Cocktail (P8340, Sigma-Aldrich), Phosphatase Inhibitor Cocktail 2 (P5726, Sigma-Aldrich), 1 mM sodium vanadate, and 1 mM phenylmethylsulfonyl fluoride (PMSF) and collected at times shown. Protein concentration was determined using the DC protein assay (Bio-Rad), measuring absorbance at 695nm in a Synergy HT or Synergy H1 Plate Reader (Biotek). Whole cell lysates were resolved by SDS-PAGE in 4-20% gradient Mini-PROTEAN TGX gels (Bio-Rad) and transferred to 0.2 μm pore size nitrocellulose membranes (Bio-Rad). Membranes were incubated for 1 h in blocking buffer (Tris-buffered saline [TBS] with 5% powdered milk), incubated with primary antibodies specific for phospho–ERK 1/2 (Thr202/Tyr204, #9101) and –MEK 1/2 (Ser217/221, clone 41G9, #9154), total ERK 1/2 (#9102) and MEK (#9122), PARP (clone 46D11, #9532), caspase 3 (clone D3R6Y, #14220), reovirus polyclonal antiserum, and *α*-tubulin (clone DM1A, #3873) overnight at 4°C. Antibodies are from Cell Signaling Technology. Membranes were washed with TBS-T (TBS with 0.1% Tween 20) and incubated with secondary antibodies conjugated to IRDye 680 or IRDye 800 (LI-COR Biosciences). Membranes were imaged using a LiCor Odyssey CLx. Images were processed and band density measured in ImageStudio (LI-COR Biosciences).

### Cell viability assay

Metabolic activity was used as a measurement of cell viability by using Presto Blue reagent (Invitrogen). MDA-MB-231 and MDA-MB-436 cells were untreated or treated with DMSO, increasing concentrations of MEK1/2 inhibitor U0126 or 25 μM pan-caspase inhibitor Q-VD-OPH for 1 h at room temperature and adsorbed with reovirus at an MOI of 500 PFU/cell for 1 h at room temperature or treated with 50 μM etoposide. Cells were washed with PBS and incubated for 0-6 days at 37°C in the absence or presence of DMSO, U0126 or Q-VD-OPH. Presto Blue (Thermo Fisher Scientific) was added at each time point for 30 min at 37°C and fluorescence (540 nm excitation/590 nm emission) was measured using black 96-well plates with clear bottom (Corning, 3904) with a Synergy HT or Synergy H1 plate reader (Biotek).

### Flow cytometric analysis of reovirus cell death

Cell viability was assessed by measuring FITC-labeled Annexin V (BioVision) (525/40 nm) and propidium iodide (690/50 nm) fluorescence using flow cytometry. MDA-MB-231 cells were pretreated with vehicle (DMSO) or 25 μM pan-caspase inhibitor Q-VD-OPH for 1 h at room temperature prior to being adsorbed with T1L, T3D, and r2Reovirus at an MOI of 500 PFU/cell for 1 h at room temperature or treated with 50 μM etoposide, washed with PBS, and incubated for 1-3 days at 37°C in the presence of DMSO or Q-VD-OPH. Cells were collected at each time point and resuspended in Annexin Cocktail (1X Annexin Buffer [10 mM HEPES, 0.14 M NaCl, and 2.5 mM CaCl_2_ in water], Annexin V-FITC, and propidium iodide). Mean fluorescence intensity (MFI) was assessed using a CytoFLEX flow cytometer (Beckman Coulter) and quantified using FlowJo software (BD Biosciences).

### Flow cytometric analysis of mitochondrial membrane potential

Mitochondrial membrane potential was measured by using tetramethylrhodamine, ethyl ester (TMRE) (Abcam). MDA-MB-231 and MDA-MB-436 cells were adsorbed with reovirus at an MOI of 500 PFU/cell or treated with 50 μM etoposide for 1 h at room temperature, washed with PBS, and incubated at 37°C for 1-3 days post-infection. Cells were stained with 100 nM TMRE at 37°C for 1 h at each time point. Cells were collected and resuspended in PBS containing 2% FBS. MFI was assessed using a CytoFLEX flow cytometer (Beckman Coulter) and quantified using FlowJo software (BD Biosciences).

### Confocal microscopy to assess cytochrome c intracellular localization

MDA-MB-231 cells plated on #1.5 glass coverslips were adsorbed with T1L, T3D or r2Reovirus at an MOI of 100 PFU/cell or treated with 50 μM etoposide for 1 h at room temperature, washed with PBS, and incubated at 37°C for 0-4 days post-infection. At each time point, cells were collected and incubated with media containing 300 nM MitoTracker Red-CMX Ros (Thermo Fisher) for 1 h at 37°C. Stained cells were fixed with 4% paraformaldehyde (PFA) in PBS for 20 min at room temperature. PFA was quenched with equal volume of 0.1 M glycine, cells were washed with PBS and stored at 4°C. Cells were treated with 0.1% Triton X100 and washed with PBS-BGT (PBS/0.5% BSA/0.1% Glycine/0.05% Tween 20), incubated with reovirus polyclonal antiserum for 1 h at room temperature, and washed with PBS-BGT. Cells were incubated with secondary antibody (Alexa 405, Thermo Fisher Scientific) and AlexaFluor488-conjugated cytochrome C monoclonal antibody (BD Pharmingen, cat. 560263), washed with PBS-BGT and mounted on coverslips with Aqua Poly/Mount (Polysciences Inc.). Cells were imaged by confocal microscopy using Olympus IX81 laser-scanning confocal microscope using a PlanApo N 60× oil objective with a 1.42 numerical aperture (NA). Pinhole size was the same for all fluorophores. Single sections of 0.44 μm thickness from a Z-stack are presented. Whole images were only adjusted for brightness and contrast.

### Measuring caspase 9 activity

Caspase 9 activity was measured by using Caspase-9 Colorimetric Assay Kit (Biovision). MDA-MB-231 cells were adsorbed with T1L, T3D, and r2Reovirus at an MOI of 500 PFU/cell for 1 h at room temperature or treated with 50 μM etoposide, washed with PBS, and incubated for 1-3 days at 37°C. Caspase 9 activity was measured at each time point using manufacturer’s instructions and reading absorbance in a clear 96-well plate (Greiner) with a Synergy HT or Synergy H1 plate reader (Biotek).

### Measuring caspase 3/7 activity

Caspase 3/7 activity was measured by using Caspase Glo reagent (Promega). MDA-MB-231 and MDA-MB-436 cells were adsorbed with reovirus at an MOI of 500 PFU/cell for 1 h at room temperature or treated with 50 μM etoposide, washed with PBS, and incubated for 1-3 days at 37°C. Alternatively, MDA-MB-231 cells were adsorbed with reovirus at an MOI of 500 PFU/cell for 1 h at room temperature, washed with PBS, incubated at 37°C, and treated with 50 μM etoposide 1 day post-infection. Caspase 3/7 activity was measured at each time point by incubating cells in equal amounts of Caspase Glo solution and cell media for 30 min at RT and reading luminescence in a white 96-well plate (Greiner) with a Synergy HT or Synergy H1 plate reader (Biotek).

### Electrophoretic mobility of reovirus

5 × 10^10^ particles of purified reovirus were mixed with 2× SDS sample buffer (20% glycerol, 100mM Tris-HCl [pH 6.8], 0.4% SDS, and 3 mg bromophenol blue) and separated by SDS-PAGE using 4-to-20% gradient polyacrylamide gels (Bio-Rad Laboratories) at 10 mAmps for 16 h. The gel was stained with 5 μg/ml ethidium bromide for 20 min and imaged using the ChemiDoc XRS+ system (Bio-Rad).

### Statistical analysis

Mean values for independent experiments were compared using one or two-way analysis of variance (ANOVA) with Tukey’s or Dunnett’s multiple-comparison test (Graph Pad Prism). *P* values of < 0.05 were considered statistically significant.

## Acknowledgements

We would like to thank Jessie Wozniak and Nicholas Harbin for careful review of the manuscript and useful suggestions. This work was supported by funding from the Children’s Healthcare of Atlanta and the Pediatric Research Institute, Winship Comprehensive Cancer Institute #IRG-14-188-01 from the American Cancer Society, and the National Institutes of Health (R01 AI146260) (B.A.M.). Flow cytometry experiments were performed in the Emory Pediatrics Flow Cytometry Core (UL1TR002378). Imaging was performed at the Emory Integrated Cellular Imaging Core (2P30 CA138292-04 and the Emory Pediatrics Institute). The funders had no role in the study design, data collection and analysis, decision to publish, or preparation of manuscript. The reassortant virus used (r2Reovirus) is part of International Patent Application No PCT/US2019/036151.

